# A revised 1.6 Å structure of the GTPase domain of the Parkinson’s disease-associated protein LRRK2 provides insights into mechanisms

**DOI:** 10.1101/676627

**Authors:** Chun-Xiang Wu, Jingling Liao, Yangshin Park, Neo C. Hoang, Victoria A. Engel, Li Wan, Misook Oh, Ruslan Sanishvili, Yuichiro Takagi, Steven M. Johnson, Mu Wang, Mark Federici, R. Jeremy Nichols, Alexandra Beilina, Xylena Reed, Mark R. Cookson, Quyen Q. Hoang

## Abstract

Leucine-rich repeat kinase 2 (LRRK2) is a large 286 kDa multi-domain protein whose mutation is a common cause of Parkinson’s disease (PD). One of the common sites of familial PD-associated mutations occurs at residue Arg-1441 in the GTPase domain of LRRK2. Previously, we reported that the PD-associated mutation R1441H impairs the catalytic activity of the GTPase domain thereby traps it in a persistently "on" state. More recently, we reported that the GTPase domain of LRRK2 exists in a dynamic dimer-monomer equilibrium where GTP binding shifts it to the monomeric conformation while GDP binding shifts it back to the dimeric state. We also reported that all of the PD-associated mutations at Arg-1441, including R1441H, R1441C, and R1441G, impair the nucleotide-dependent dimer-monomer conformational dynamics of the GTPase domain. However, the mechanism of this nucleotide-dependent conformational dynamics and how it is impaired by the mutations at residue Arg-1441 remained unclear. Here, we report a 1.6 Å crystal structure of the GTPase domain of LRRK2. Our structure has revealed a dynamic switch region that can be differentially regulated by GTP and GDP binding. This nucleotide-dependent regulation is impaired when residue Arg-1441 is substituted with the PD-associated mutations due to the loss of its exquisite interactions consisting of two hydrogen bonds and a π-stacking interaction at the dimer interface.

**Significance Statement:** Mutations in LRRK2 are associated with familial Parkinson’s disease, so understanding its mechanism of actions and how they are changed by the disease-associated mutations is important for developing therapeutic strategies. This paper describes an atomic structure of the G-domain of LRRK2 revealing that the conformational dynamics of the switch regions are potentially important for its normal function. It further shows that a disease-associated mutation could lock the G domain in a persistently active-like conformation, thus perturbing its normal function.

---

Mutation in the Leucine-Rich Repeat Kinase 2 (LRRK2) is a common cause of familial Parkinson’s disease (PD)(1–5). LRRK2 is a large (2,527 amino acids) multi-domain enzyme consisting of 7 putative domains(2), including a HEAT repeat, an Ankyrin repeat, a Leucine-rich repeat (LRR), a Ras-like GTPase domain called ‘Ras of complex proteins’ (ROC), a unique domain called C-terminal of ROC (COR), a kinase domain (Kin), and a C-terminal WD40 repeat(6). The most common PD-associated mutations (G2019S and R1441G/C/H) are found in the central three domains, ROC-COR-Kin, that constitutes the catalytic core of the protein. Residue Gly-2019 is predicted to reside in the activation loop of the kinase domain, and the G2019S mutant has been shown to have higher kinase activity than that of the wild-type enzyme(7, 8). Thus, the glycine to serine substitution mutation, which introduces a hydroxyl group, potentially mimics the activated state of the kinase.

The effects of the PD-associated mutations in the ROC domain at residue Arg-1441 (R1441G/C/H) have been less understood. Several studies have shown that GTP binding to the ROC domain up regulates LRRK2 kinase activity(9, 10) and, conversely, inhibition of ROC decreases kinase activity(9–13), thus indicating that the ROC domain could modulate the activity of the kinase domain. Moreover, the PD-associated mutations at Arg-1441 have been shown to also result in a higher LRRK2 kinase activity than that of the wild-type(8, 9, 12), suggesting that residue Arg-1441 is vital for the normal functioning of ROC. Indeed, we recently engineered a stable construct of ROC (ROC_ext_) and showed that the R1441H mutation perturbs the hydrolytic conversion of GTP to GDP, thereby prolonging the GTP-bound "on" state of ROC(14). Subsequently, we showed that GTP-binding drives dimeric ROC_ext_ disassembly into monomers and GDP-binding oppositely shifts the equilibrium back to dimers(15). Moreover, we showed that all the PD-associated mutations at Arg-1441, as well as N1437H, impair the dimer-monomer dynamics of ROC_ext_(15, 16).

The mystery remains as to how residue Arg-1441 affects the catalytic activity of ROC and its nucleotide-dependent conformational dynamics given that the position 1441 resides outside of the regions known to be involved in the activity of GTPases, including the nucleotide-binding site, active-site, and the switch regions. Here, we show a high-resolution crystal structure of ROC_ext_ revealing the role of residue R1441 in the stability of the dimeric conformation and providing insights into how the Parkinson’s disease-associated mutations R1441G/C/H impair its conformational dynamics.

## Results and Discussion

### The architecture of the ROC domain of LRRK2

While a structure of the ROC domain of LRRK2 has been determined a decade ago (PDB ID: 2ZEJ)(17), that structure has been inconsistent with subsequent biochemical and biophysical observations. For example, the 2ZEJ structure showed an obligate dimer with an active-site composed of one half from each protomer; however, biochemical studies have demonstrated that ROC dynamically interconverts between its dimeric and monomeric conformations and that the monomers are catalytically active(14, 15). Thus, the examination of the 2ZEJ structure did not provide us with the insights that we sought to understand the conformational dynamics ROC and the mechanisms of the PD-associated mutations. We attempted to glean these insights from structures of the homologous protein Roco from *C. tepidum*(18, 19); however, we found significant differences between the ROC domain of human LRRK2 (HsROC) and that of *C. tepidum* Roco (CtROC). For example, the PD-associated residues in HsROC are not conserved in CtROC, including residue R1441 in HsROC; therefore, the potential biochemical and structural effects of the PD-associated mutations cannot be inferred directly from the CtROC structures. Due to these reasons, we set out to determine the structure of the human ROC_ext_ construct to examine the structural bases for the biochemical and conformational dynamics we observed(14, 15, 20).

We have determined a crystal structure of ROC_ext_ to 1.6 Å resolution using X-ray crystallography. The initial electron density map was calculated using a combination of phases determined with SHARP(21) using single-wavelength anomalous dispersion of selenomethionine (SeM-SAD) from crystals of selenomethionine-substituted proteins, which diffracted to 3.7 Å, and those calculated by the molecular replacement (MR) method using the program Phenix(22) with a 1.9 Å native dataset. To obtain a better crystal quality and higher resolution, we performed “surface engineering” by substituting two surface lysine residues for alanine (K1460A and K1463A). The surface engineered mutant (KA) retained wild-type activity and structure (Fig. S1). The data from crystals of the surface-engineered protein extended to 1.6 Å resolution and were used for model building and refinement. A contiguous polypeptide chain ranging from residues 1331 to 1518, which nearly encompassed the entire ROC_ext_ construct (1329-1520), was unambiguously fitted into the electron density and refined to convergence with R-factor and R-free of 14% and 16%, respectively (Sup. Table 1).

The asymmetric unit (AU) consists of a homodimeric structure of ROC_ext_ related by 2-fold symmetry. The interactions at the dimer interface are extensive, burying 6461 Å^2^ of surface area (Fig. 1a-c). The intermolecular interactions are mediated chiefly by the structures that are known as Switch I, Switch II, and the InterSwitch, herein collectively referred to as the Switch regions (Fig. 1b-e). The dimeric interface is unusual in that the InterSwitch is flipped open and inserted into its dimeric partner *in trans* with both monomers wrapped around each other like a pretzel (Fig. 1a). This dimeric conformation has been previously observed in *M. Musculus* Rab27b, which showed a similar extended InterSwitch structure that reaches across the homodimeric interface(23). Apo Rab27b exists in solution as a monomer with its InterSwitch extended, however, upon binding to its effector Slac2-a it folds into a typical G-protein conformation with a retracted InterSwitch and adopting a conformation similar to that of the typical Ras-family of G-proteins(24).

**Figure 1.**
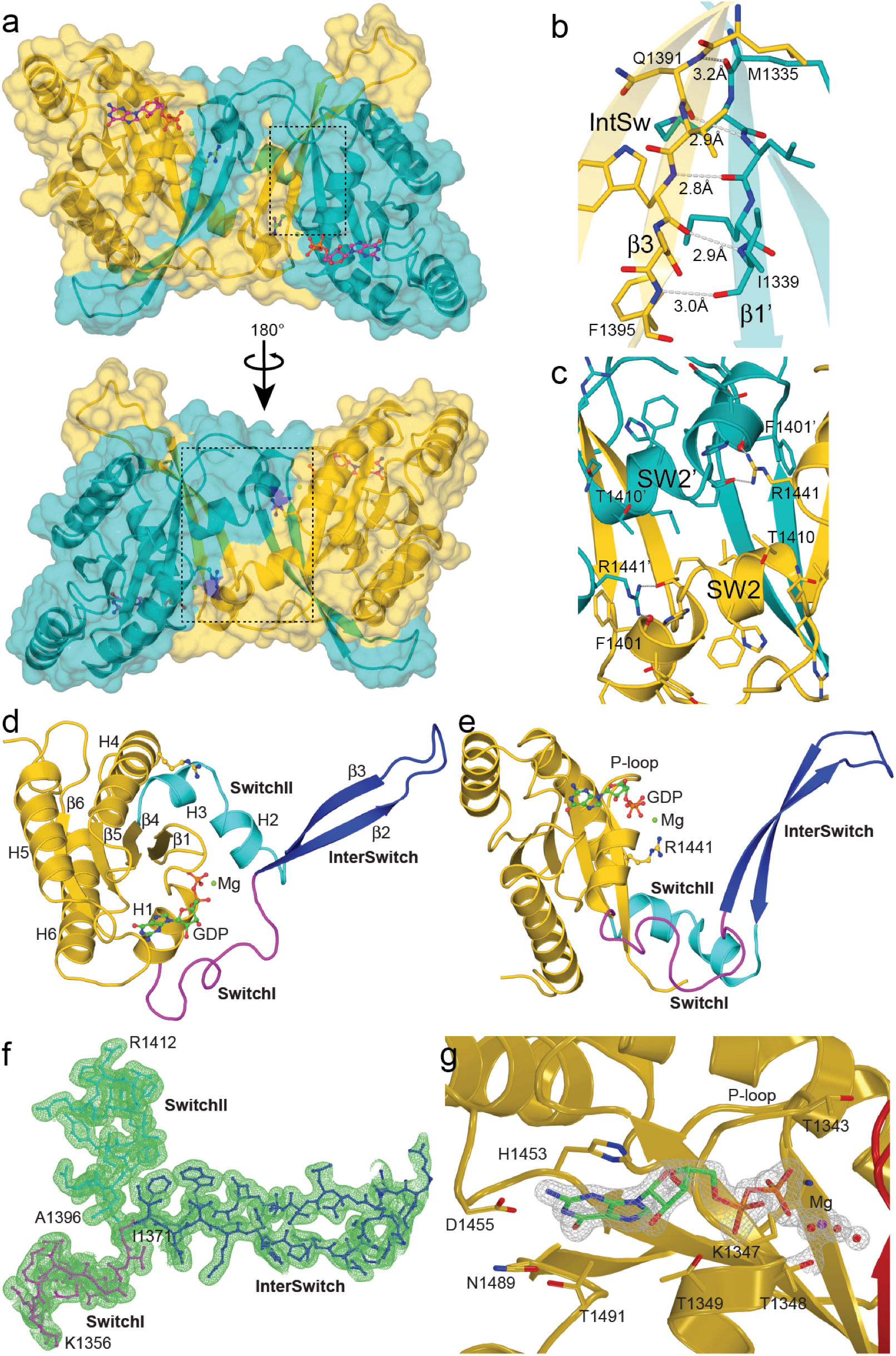
Structure of ROC_ext_. **a**) Crystal structure of ROC dimer (1.6 A) shown in semi-transparent surface and ribbon presentations. Chain A is colored orange and chain B colored teal. The dotted lines outline the areas presented in **b**) and **c**). **b**) Enlarged view of the area highlighted in top panel of **a**) showing extensive hydrogen bonding at the dimer interface. **c**) Enlarged view of the area highlighted in bottom panel of **a**) showing a hydrophobic patch at the dimer interface, which is capped at both ends by residue R1441. **d**) and **e**) ribbon presentation showing signature structural features of a small GTPase, including Switch I (magenta), InterSwitch (blue), and Switch II (teal). GDP is shown as rod-and-ball model. **f**) and **g**) electron-density map (2FoFc) of the Switch regions and GDP, respectively, contoured at 1.0 σ.

The tertiary structure of ROC_ext_ is very similar to other G-proteins in that it consists of all the characteristic structural elements, including Switch I, Switch II, P-loop, and the G-binding motifs (Fig. 1d-g). The Switch I and Switch II regions of ROC_ext_ are distal from the GDP-binding site, as they can also occur in GDP-bound Ras-like GTPases. However, in contrast to Ras, the InterSwitch of ROC_ext_, which is flanked by the Switch I and Switch II, is flipped outward and protrude away from the protein core - like a ‘thumbs-up’ hand gesture (Fig. 1d,e). A well-defined molecule of GDP is bound in a location similar to the nucleotide-binding site in Ras (Fig. 1g).

### Comparison of ROC_ext_ with the previously reported ROC structure

Superposition of the structure of ROC_ext_ with 2ZEJ revealed two significantly different structures (RMSD = 10.4 A^2^). A large portion of the N-terminus of 2ZEJ (residues 1335-1397) consisting of β1, H1, β2, and β3 are flipped outward and detached from the rest of the protein, in effect, breaking the G-domain into two distinct halves with the N- and C-terminus separated from each other (Fig. 2a). Removing the non-overlapping moieties, β1 and H1, from the structural superposition resulted in two structures that are nearly identical (RMSD = 0.4 A^2^). Close inspection revealed that the β1 and H1 moieties in chain A of 2ZEJ occupy precisely the same locations of the β1 and H1 of chain B in our ROC_ext_ structure (Fig. 2b), thus suggesting that the structural differences might be simply due to the difference in chain assignment of β1 and H1. Indeed, superposition of the two structures as dimers showed nearly identical structures (RMSD = 0.3 A^2^). Inspection of the electron-density map of the 2ZEJ structure revealed that the junctions between H1 and β2, where the Switch I would reside, lacked interpretable density (Fig. S2, S3); thus rendering the task of assigning the chain ID for the H1 and β2 segment ambiguous. Since our electron-density map unambiguously defined a contiguous polypeptide chain, we propose that the ROCext presented herein represents a revised structural model. Indeed, when the chain ID assignments for β1 and H1 in 2ZEJ were swapped (A to B and vice versa), the resulting structure looked nearly identical to our ROC_ext_ (Fig. S4).

**Figure 2.**
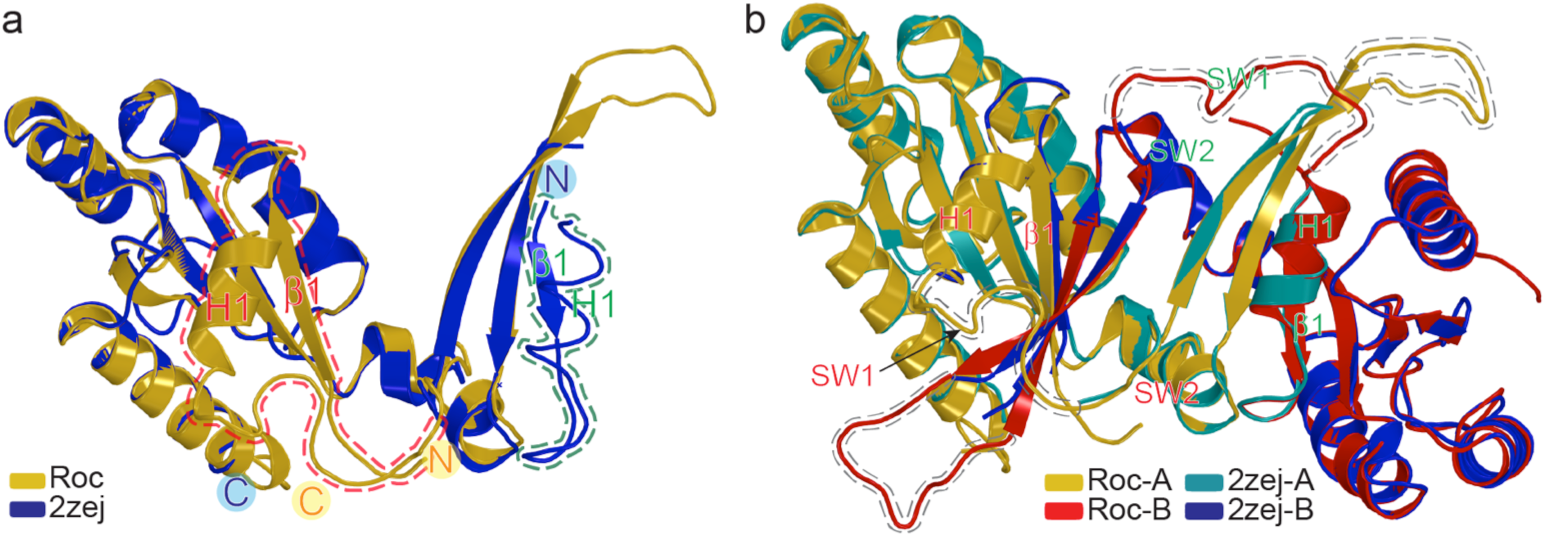
Superposition of ROC_ext_ with 2ZEJ. **a**) Structure of ROC_ext_ (gold) superimposed with the ROC structure in the literature, 2ZEJ, (blue). The equivalent structural elements that are located in different locations in the two structures are outlined with dashed lines. **b**) Superposition of ROCext dimer (gold and red) with 2ZEJ (teal and blue), showing overlap of all atomic positions of 2ZEJ with those of ROC_ext_. Structural elements in ROC_ext_ missing in 2ZEJ are highlighted with grey dashed lines.

### Comparison of human ROC with *C. tepidum* Roco

Because of the difficulty in obtaining sufficient quantity of LRRK2 amenable for detailed investigations, insights into its structure and function have been inferred from the amoeba and bacterial homologs(18, 25). Notably, much insight into the ROC domain of LRRK2 (HsROC) has been inferred from a ROC structure of the bacterium *C. tepidum* Roco protein (CtROC) (PDB ID: 3DPU, 6HLU). For example, the mechanism of GTPase hydrolysis of HsROC has been proposed based on the studies of CtROC, where the activation occurs upon an arginine finger exchange in the active-site mediated by homodimerization (known as the GTPase activation by dimerization model or GAD) (18),(26). However, we have recently shown that ROC_ext_ is catalytically active as a monomer, suggesting that the mechanism of HsROC might be different from that of CtROC. To assess the similarity and differences between CtROC and ROC_ext_, we superimposed the two structures. Although they share low sequence identity (25%), the overall architecture of the two structures are remarkably similar (RMSD = 2.2 Å^2^) (Fig. 3a). Nevertheless, the superimposed structures did reveal several important differences: firstly, the ROC-ROC dimeric interface of CtROC contains the P-loop, whereas the dimeric interface of ROC_ext_ comprises of the Switch regions (Fig. 3b). Note that neither the dimerization mode of CtROC nor ROC_ext_ supports a GAD mechanism of GTPase activation, as there is no arginine finger-exchange occurring in the active sites. Second, the disease-associated residues R1441, N1437, and R1398 in HsROC structurally aligned with residues Y558, H554, and Q519 of CtROC, respectively. The difference in amino acids at these positions is important in that the substitution of residue Arg1441 in HsROC for Tyr found in the equivalent position in CtROC, and residue N1437 in HsROC for His of CtROC both resulted in impairment of the conformational dynamics of ROC_ext_(15, 16) (Fig. 3c). Finally, the arginine residue R543 of CtROC that is proposed to serve as the GTPase-activating arginine finger is structurally equivalent to residue S1422 of HsROC (Fig. 3d). While the studies of CtRoco have been instrumental for the understanding of the overall architecture of ROC-COR, these structural differences suggest caution in deducing the structure and mechanisms of HsROC from the observations of CtROC.

**Figure 3.**
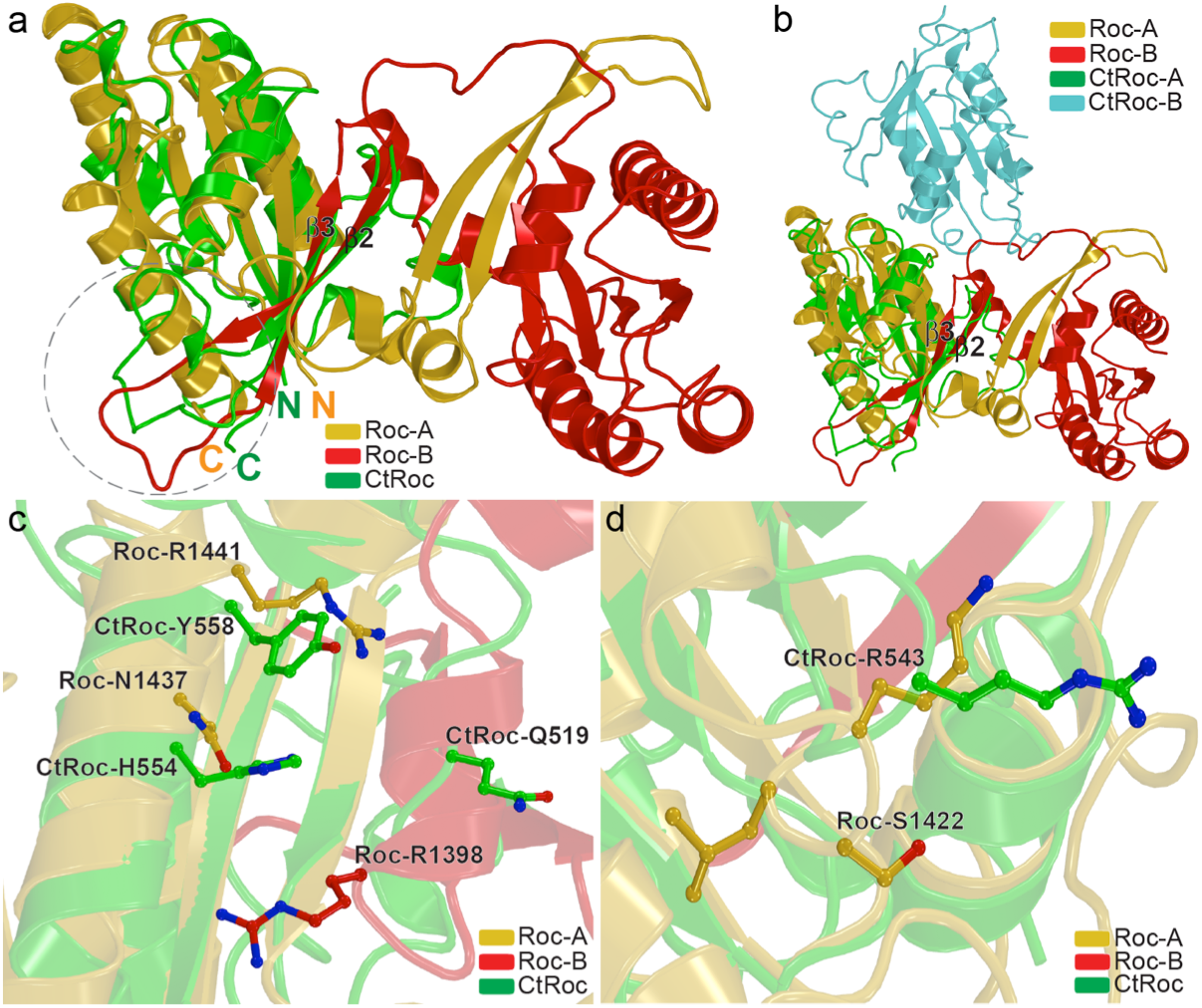
Comparison with the structure of CtROC. **a**) Superposition of ROC_ext_ dimer (gold and red) with CtROC (green), showing that the two structures overlap well, other than the turn/loop of the InterSwitch between b1 and b2 (highlighted in grey dashed line). **b**) Superposition of A-chain of ROC_ext_ with A-chain of CtROC (6HLU), showing that the two structures form dimers via different interactions and surfaces. **c**) and **d**) Enlarged view of the superposition of ROC_ext_ with CtROC, showing the amino acid differences between the structures.

### The crystal structure of ROC_ext_ represents its in-solution dimeric conformation

As reported previously, ROC_ext_ exists in solution in a dynamic monomer-dimer equilibrium. To investigate whether the dimer in solution is the same as the dimer in crystal, we introduced an intermolecular disulfide bond between two residues that are in close proximity of each other, but each residing on opposite side of the dimer interface of the crystal dimer. This disulfide bond links *in trans* the helix H2 of one protomer to the helix H4 of the other protomer (residues R1398C and W1434C) (Fig. 4a, Fig. S5a, b).

**Figure 4.**
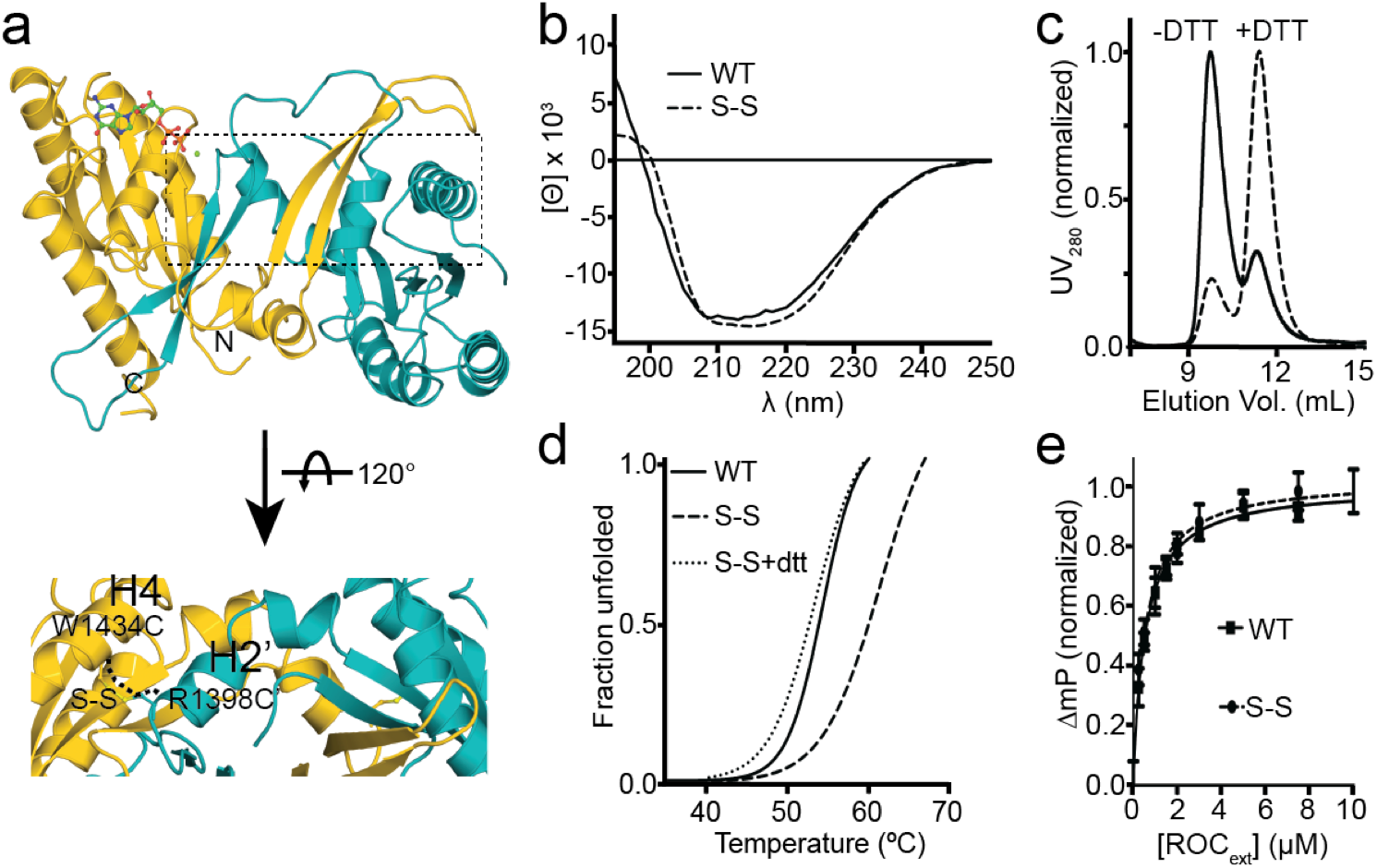
Engineered disulfide-stabilized ROC dimer. **a**) A calculated model of ROC dimer consisting of an engineered disulfide bond between residues 1398 and 1431 (S-S). **b**) CD spectrum showing similar profile between the S-S dimer and wild-type. **c**) Size-exclusion chromatography elution profiles showing that the S-S dimer has lost the ability to convert to dimer and then regains that capability when the cysteine residues are reduced with DTT. **d**) Thermofluor denaturation assay showing that the S-S dimer is more thermally stable than the wild-type, but reverts to wild-type melting temperature when reduced with DTT. **e**) Fluorescence polarization assay showing that the S-S dimer binds GDP with a similar affinity to wild-type.

The resulting disulfide-stabilized construct (S-S) formed both monomeric and dimeric conformations reminiscent of the WT (Fig. S6 & S7). Circular dichroism (CD) spectroscopy showed no significant structural differences between the S-S mutant and the WT dimers (Fig. 4b). However, the S-S dimer loses the ability to convert to monomers upon nucleotide binding under disulfide bond-stabilizing oxidative conditions, but in disulfide bond-breaking reducing conditions, its propensity to form monomers is restored to the wild-type level (Fig. 4c). Likewise, the S-S dimer is more thermostable than the WT under oxidizing conditions, but returned to WT Tm upon reduction with DTT (Fig. 4d). Finally, the GDP-binding affinity of the S-S dimer is the same as that of the WT (Fig 4e).

Taken together, these results indicate that the *in crystallo* dimer is structurally the same as the dimer we observed in solution, and therefore, insights gleaned from the crystal structure would be reflective of the structure in solution.

### Insights into the mechanism of ROC

Understanding the mechanism of the ROC domain of LRRK2 is important for understanding its overall function and for the development of mechanism-based therapeutic strategies. However, the current understanding of ROC has been mostly inferred from other well-characterized G-proteins. They all function as molecular switches that toggle between "on" and "off" upon binding to GTP or GDP, respectively. The binding of GTP typically causes the stabilization of two flexible structures, known as Switch I and Switch II, as they both interact electrostatically with the γ-phosphate of GTP. The ordering of the switch regions presents a surface for interacting with downstream effector proteins, and its disordering upon GTP hydrolysis (or exchanged for GDP) resets the molecular switch back to the "off" state(27–29). Indeed, the ROC domain of LRRK2 has been shown to possess some of these characteristics, including guanine nucleotide binding, GTP hydrolysis, and nucleotide-dependent conformational changes. However, ROC has also been shown to present several characteristics that do not fit precisely into the typical small G-protein model. For example, ROC always exist in tandem with a domain called C-terminal of ROC (COR) within a large protein, ROC remains active event when mutated to residues that would abolish the activity of the other Ras-family G-proteins, and its ‘ability’ to hydrolyze GTP is required for its activity but not the hydrolysis event itself per se(30). Although it is evident that the ROC domain of LRRK2 is a bona fide G-protein, these differences suggest that the mechanism of ROC might be different from that of the typical small G-proteins.

To gain insights into the mechanism of ROC, we examined the atomic structure of ROC_ext_ to glean the structural bases for its biochemical activities. We recently reported that ROC_ext_ exists in a dynamic monomer-dimer equilibrium and that GTP or GDP binding drives monomerization or dimerization, respectively(15). The structure of ROC_ext_ reveals that the dimeric configuration of ROC_ext_ is predominantly stabilized by five hydrogen bonds between β3 in the InterSwitch of one protomer and β1 in the core of the other and arranged parallel with respect to each other like a zipper (Fig 1b,c). The labile nature of these hydrogen bonds is well suited to provide the plasticity necessary for the ‘zipping’ and ‘unzipping’ in the interconversion between the dimeric and monomeric conformations.

Since we have only the GDP-bound structure, how the binding of GTP would drive the unzipping of the dimer is unclear. However, close examination of our ROC_ext_ structure revealed plausible clues. The residues flanking either side of the InterSwitch possess dihedral angles that are conformationally strained (ϕ, ψ angles of S1360 and Q1411 are −93°, −90° and −105°, 96°, respectively) and, as such, the stored energy in these hinge regions could potentially act as a set of spring-loaded hinges that provide the driving force for unzipping the dimer triggered by GTP binding. This led us to question how does GTP binding trigger the process of monomerization. The mechanism of GTP-induced conformational changes in small G-proteins involves the γ-phosphate group of GTP interacting with a threonine residue of Switch I and a glycine residue of Switch II. To investigate whether or not this occurs in ROC_ext_, we examined the structure of ROC_ext_ to see if it consists of the essential γ-phosphate-interacting threonine and glycine. To do so, we built a homology model of GTP-bound conformation of ROC_ext_ by using a GTP-bound conformation of Ras (PDB ID: 6Q21) as a modeling template for the switch regions. Superposition of 6Q21 with the GTP-bound homology model of ROC_ext_ reveals that it does indeed consist of a threonine residue (T1368) in Switch I and a glycine residue (G1397) in Switch II located at practically the same locations as the γ-phosphate-interacting residues, T35 and G60, respectively, of Ras (Fig. 5a,b and Fig. S8). The conservation of T1368 and G1397, both in primary and tertiary structures, with the corresponding γ-phosphate-interacting residues of Ras suggests that their interaction with the γ-phosphate of GTP could analogously trigger conformational changes of the Switch regions. We noted that GTP-binding to Ras triggers the ordering of the Switch I and Switch II; however, the structure of ROC_ext_ suggests that conformational changes in the two switches also necessitate a conformational change of the InterSwitch. This opens the possibility that the conformational changes of the InterSwitch might be involved in yet to be identified interactions analogous to Rab27b interaction with Slac2a(23, 24).

**Figure 5.**
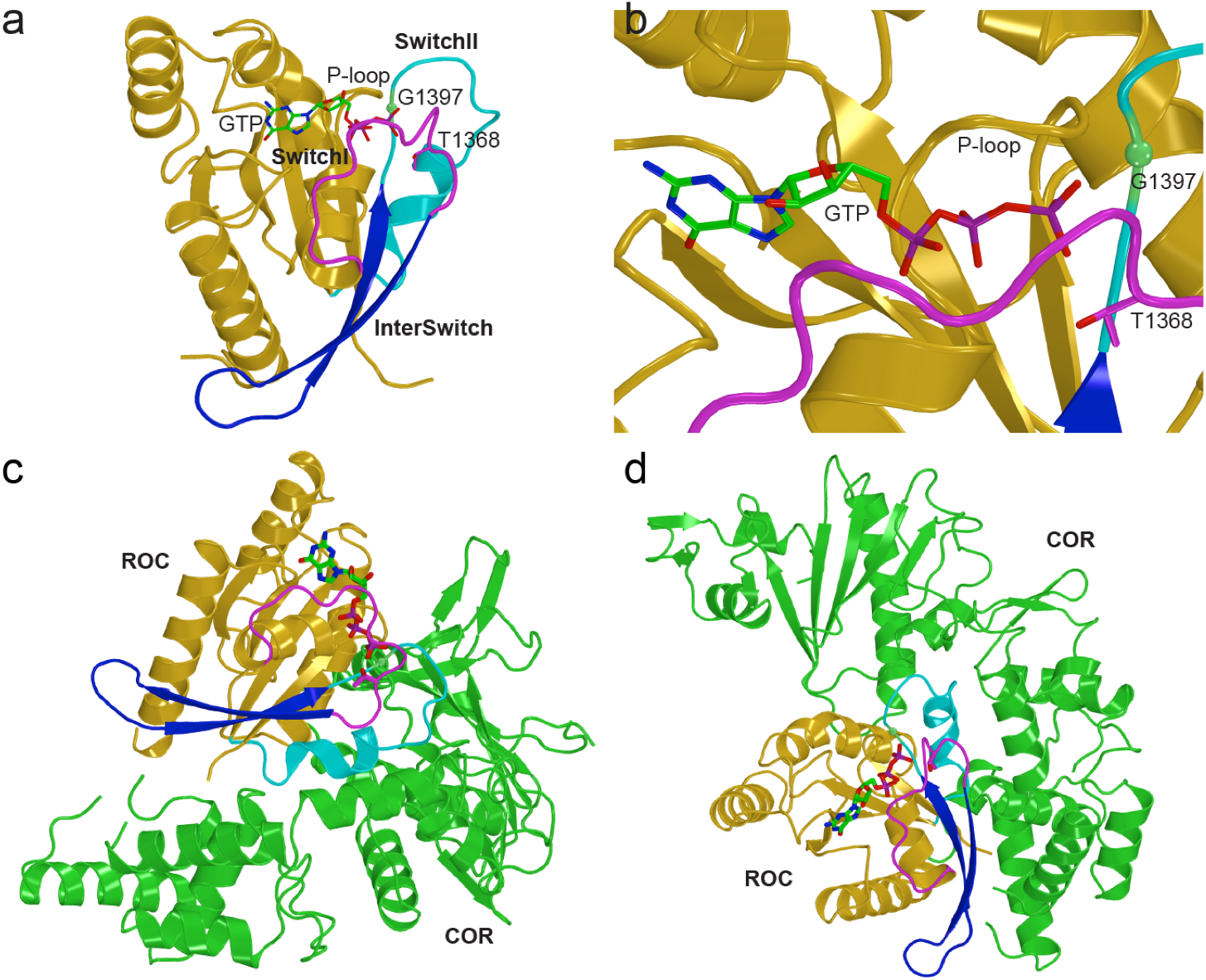
Modeling of a monomeric form of ROC. **a**) and **b**) A model of monomeric GTP-bound ROC built based on the ROC_ext_ dimer and the switch regions of the structure of GppNHp-bound Ras (6q21), showing that ROC has all the structural elements and residues that are involved in the GTP-mediated conformational changes in Ras. **c**) and **d**) docking of GTP-bound ROC model to the COR domain of CtRoco (6HLU), showing the surface complementarity between the two structures.

Biochemical and structural data, shown herein and in the literature, have sufficiently demonstrated that the ROC domain of LRRK2 forms an independently folded and functional unit; however, how it interacts intramolecularly with the other domains, especially the adjacent COR domain, remains unclear. To investigate this, we docked the dimeric form of ROC as found in our crystal structure, the monomer of our crystal structure, and the GTP-bound model of ROC (ROC-GTP) mentioned above, to the COR domain of the CtRoco protein (PDB ID: 6HLU)(18, 19). The crystal structure of ROC_ext_ showed significant clashes with the docked CtRoco COR domain. Even in the monomeric form, the Switch regions clashed with the CtRoco COR domain. However, the ROC-GTP monomeric model docked to the CtRoco COR domain well with no backbone clashes between the two domains, and the surface containing the Switch regions, especially the InterSwitch, interfaces with the COR domain with remarkable complementarity (Fig. 5c,d).

Taken together, the data suggest that the Switch regions of the ROC domain of LRRK2 interact directly with its COR domain and that the nucleotide-dependent conformational changes of the Switch regions might modulate this interaction. However, more work is needed to validate the insights gleaned from the ROC-COR docking exercise.

### Insights into the mechanism of disease-associated mutations

Several lines of evidence have shown that LRRK2 harboring the disease-associated mutation R1441C/G caused a reduction in GTPase activity(12, 14, 31-33) and GTP binding to the ROC domain modulates kinase activity(10, 13, 30). These data suggest that the GTPase activity of LRRK2 potentially regulates its kinase activity. Recently, we reported that, in addition to a reduction in GTPase activity, a common biochemical consequence of all three PD-associated mutant constructs, including R1441H, R1441G, and R1441C, is a loss of the monomer-dimer conformational dynamics that we observed in the wild-type protein(15).

To gain insights into how the PD-associated mutants led to a loss of ROC conformational dynamics, we examined the crystal structure of ROC_ext_. The structure revealed that residue R1441 reaches across the dimer interface and interacts exquisitely with the opposing protomer (Fig. 1 and Fig. 6). The interaction consists of an arginine-π-stacking between R1441 and F1401. The strong π-stacking interaction with the phenyl ring of F1401 orients the plane of the R1441 guanidinium moiety such that its two ω-amines are ideally positioned for hydrogen-bonding with the backbone carbonyl oxygen of F1401 and sidechain hydroxyl of T1404 (Fig. 6b). This Arg-Phe π-stacking interaction is typically found in proteins where strong and malleable interactions occur(34), thus suggesting that residue R1441 might be essential for the stability and plasticity of ROC. In addition to the π-stacking, the sidechain of R1441 fits into the pocket between residues F1401 and T1404 with perfect Van der Waals complementarity (Fig. S9). Given the intricacy of the interactions and the tight complementarity involved, it appeared that no other natural amino acid would adequately substitute for the arginine at position 1441. To test this, we substituted R1441 for a relatively conserved amino acid, lysine, which preserved a positively charged amine and a long aliphatic sidechain. We found that the R1441K mutant adopted only the monomeric conformation and its activity was significantly reduced compared to that of the WT (Fig. 6d), suggesting that the intricate interactions conferred by the sidechain of residue R1441 is critical for maintaining a monomer-dimer equilibrium. This is also consistent with our recent report that substituting arginine1441 for a tyrosine resulted in a constitutive dimer(15). Now that the structure is available, we can see that a Tyr residue at position 1441 could form one hydrogen bond and a strong π-π-stacking interaction with phenylalanine 1401, which would be much less malleable than that of the WT.

**Figure 6.**
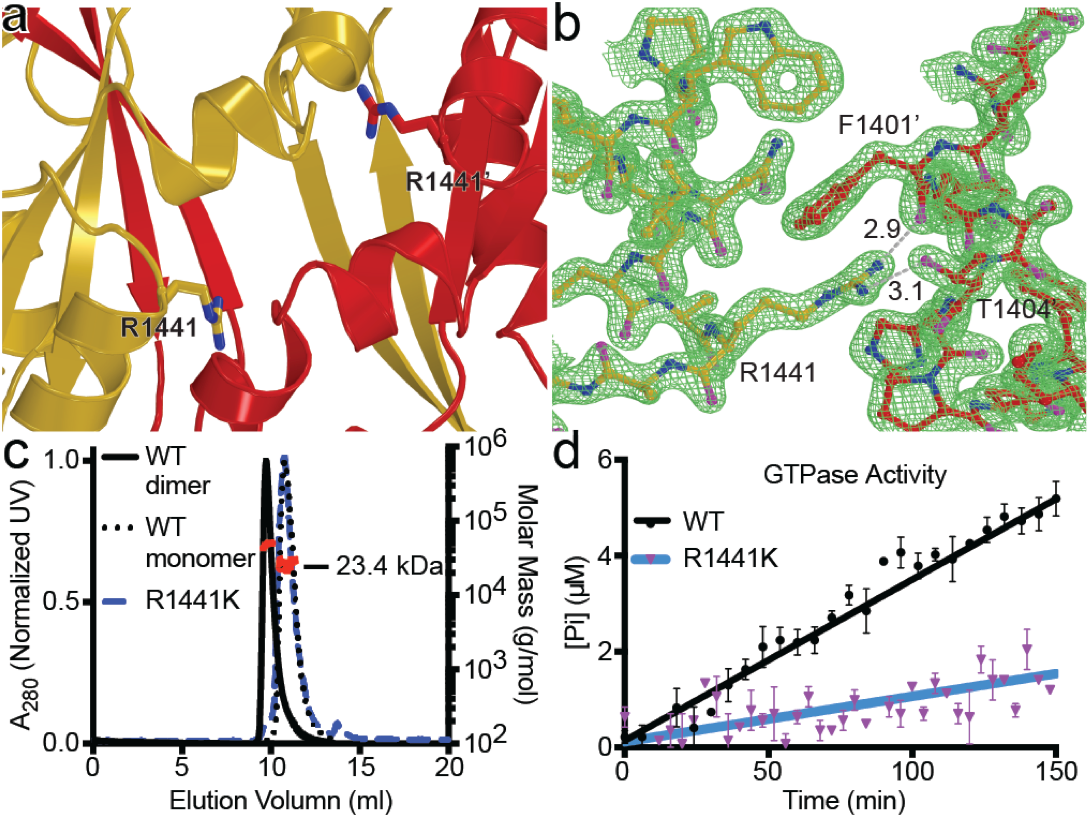
Role of R1441 residue in conformational dynamics of ROC. **a**) and **b**) Enlarged view of the dimeric interface of ROC_ext_ around residue R1441 (rod-bond model). The electrondensity (2FoFc at 1s) is shown as green mesh. **c**) SEC-MALS of the R1441K mutant form of ROC (blue line) compared to the WT dimer (black line) and WT monomer (black dotted line), showing that substituting arginine for lysine at position 1441 completely abolished dimerization. **d**) GTPase activity of R1441K (purple and blue) compared to WT (black), showing that the R1441K mutation impaired GTPase activity.

Taken together, the structure is consistent with all the biochemical phenomena we observed in solution and further supports that R1441 is uniquely required for the normal nucleotide-dependent conformational dynamics of ROC_ext_, and suggests that the PD-associated mutations at residue R1441 impair this process by disrupting the exquisite interactions endowed by the guanidinium moiety of the arginine residue.

## Conclusion

Previous knowledge of the structure and function of the ROC domain of LRRK2 have been primarily inferred from a homologous protein CtRoco due to a lack of LRRK2 protein samples amenable for detailed biochemical and biophysical studies. At the onset of our investigation into the structure and function of ROC, we were uncertain whether the domain in isolation and in the absence of the other domains would fold and function independently. However, we now know that it does, in both accounts, which has enabled us to carry out detailed biochemical and biophysical studies(14–16), as well as herein described, to determine its atomic structure. Our studies unambiguously demonstrated that ROC_ext_ forms an independently folded domain that consists of structural features, nucleotide binding properties, and enzymatic activity that are consistent with all the hallmarks of a Ras-like G-protein. It is noteworthy that the structure shows the N- and C-terminus of ROC_ext_ are in proximity of each other, as they occur in the other Ras-family G-proteins, and that they would be unperturbed by the conformational changes that occur in the processes of nucleotide exchange, hydrolysis activity, and monomer-dimer interconversions; therefore, should these activities occur in the full-length LRRK2, they can do so without requiring the movement of the flanking domains.

Although it is theoretically possible for ROC to form a homodimer in the context of the full-length LRRK2, it remains unclear whether or not that occurs, as there is insufficient data to make the determination. On the one hand, electron microscopy (EM) showed that full-length LRRK2 is dimeric and that the ROC domains are in close proximity for ROC-ROC interactions (although the dimeric interface is different from our structure) (35). On the other hand, size-exclusion chromatography of a ROC-COR tandem construct showed a dimeric structure, but a C-terminally truncated ROC-COR domain was mainly monomeric(36). However, the nucleotide status of these studies was unclear, and given the fact that ROC undergoes nucleotide-dependent conformational changes, the question of whether ROC forms a homodimer in the full-length LRRK2 remains to be elucidated. Given the fact that LRRK2 is a large protein it is likely that, in addition to ROC, the other domains are also involved in the dimerization of LRRK2. For example, the WD40 domain of LRRK2 forms a strong homodimer(37).

Based on the fact that the crystal structure showed that GDP-bound ROC_ext_ is homo-dimeric, SEC-MALS showed that GTP-bound ROC_ext_ is monomeric(15), and the docking models showed that ROC might interact with COR via the same surface in the homodimer interface, we imagined that homo-dimerization of the GDP-bound ROC would exclude its interaction with the COR domain and that its monomerization upon GTP binding might expose the surfaces available for hetero-dimerization of the two domains (Fig. 7).

**Figure 7.**
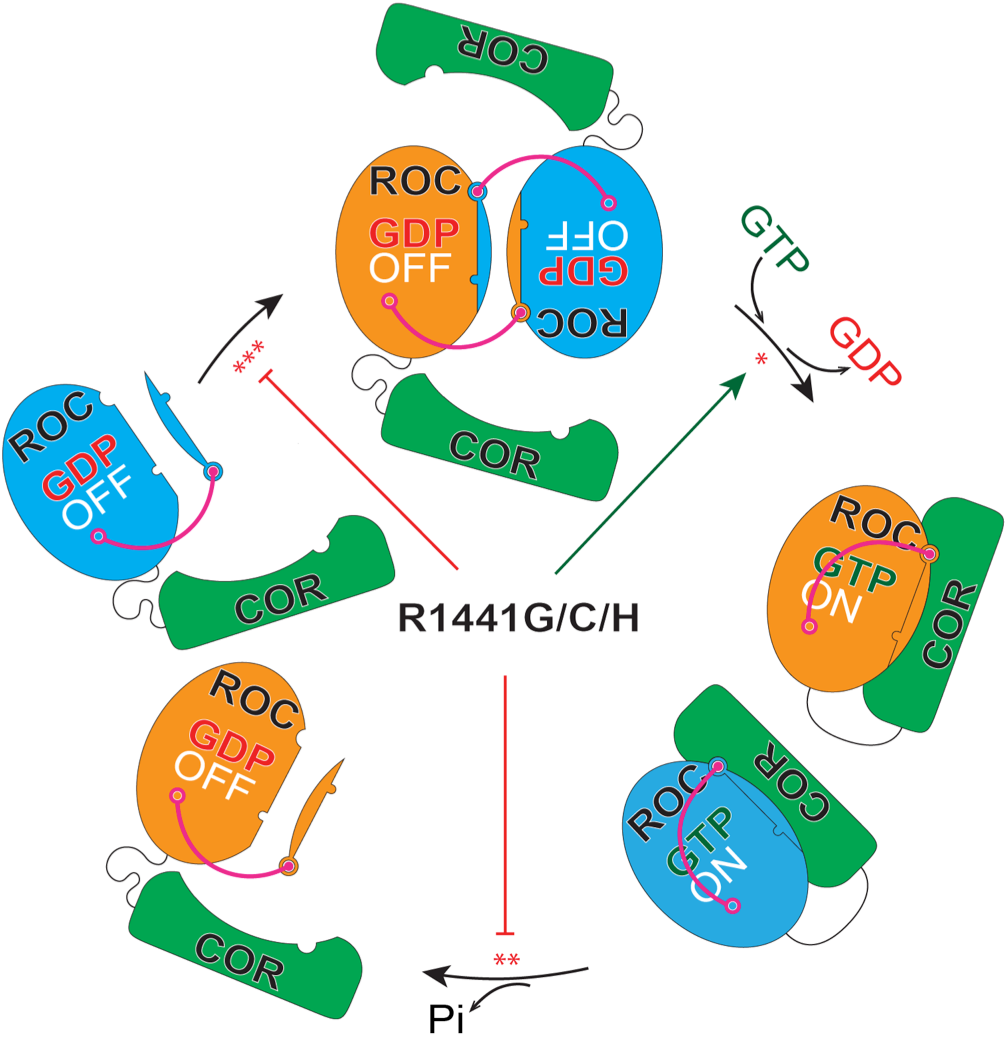
Model of nucleotide-dependent conformational changes of ROC. GDP and GTP binding regulates the dynamic interconversions between dimeric and monomeric conformation of ROC, respectively. Homo-dimerization of ROC excludes its potential interaction with the COR domain and vise versa. Mutations at residue R1441 impair this nucleotide-dependent conformational changes and thereby trapping ROC in the monomeric “on” state and potentially bound to the COR domain. *R1441H has higher affinity for GTP compared to WT. **R1441G/C/H impair GTP hydrolysis. ***R1441G/C/H lost the ability to form homo-dimers.

## Methods

A more detailed description of the methods is available in the supplemental information.

### Protein Expression and Purification

ROC_ext_ consisting of residues 1329-1520 was subcloned into a pETDuet-1 vector, expressed in Rosetta2 (DE3) *E. coli*, and purified sequentially with Ni-NTA agarose and Superdex 200 column. The disulfide bond-stabilized ROC dimer (S-S) was expressed in SHuffle T7 Express *lysY E. coli* and purified as described for the wild-type.

### Expression and Purification of Selenomethionine Substituted Protein

SeMet ROC_ext_ was expressed from Rosetta2 (DE3) *E. coli* grown in M9 minimal medium containing of a methionine synthesis-inhibiting cocktail (100 mg of Lys, Phe, Thr (Sigma Aldrich) and 50 mg of Ile, Leu, Val (Sigma Aldrich)). Expression and purification of the SeMet ROC_ext_ as described for wild-type ROC_ext_.

### Size-Exclusion Chromatography Coupled with Multi-angle Light Scattering (SEC-MALS)

Absolute molecular weight of Roc_ext_ in solution we determined by multiple-angle light scattering coupled to size-exclusion chromatography (SEC-MALS). The setup composes of an AKTA FPLC with a size-exclusion column (WTC-030S5, Wyatt Technology Corporation), a refractive index detector (Optilab T-rEX, Wyatt Tech.), and a multiple light scattering detector (Dawn HeleosII, Wyatt Tech.). Data processing and analysis were performed using the ASTRA software (Wyatt Tech.)

### Circular Dichroism Spectroscopy

CD spectra were collected on a Biologic Science Instruments MOS450 AF/CD spectrometer with a slit width of 1.0 mm and data acquisition of 1.0 s.

### Fluorescence Polarization Nucleotide-binding Assay

Binding affinity of guanine nucleotides BODIPY-FL-GTPgammaS or BODIPY-FL-GDP (Molecular Probes) were determined using fluorescence polarization measured by an EnVison 2102 Multilabel Plate Reader (Perkin Elmer).

### GTPase Activity Assay

GTPase activity of ROC_ext_ was assessed by using the Enzcheck assay kit (Invitrogen).

### Thermofluor Assay

Denaturation curves were obtained using a Sypro Orange-based method. Samples were heated in the Real-Time PCR Detection System (Mastercycler realplex, Eppendorf) from 20 to 85 ℃ and fluorescence (550 nm) recorded in increments of 0.4 ℃.

### Homology Modeling and Molecular Simulation

Homology models of the monomeric ROC domain were built based on the structures of Ras and *C. tepidium* Roco (PDB ID: 4Q21, 6Q21, 3DPU, and 6HLU) by using the program Modeller 9.19 (Andrej Sali, UCSF). Molecular simulation of the S-S mutant was performed using CHARMM(38) and NAMD(39).

### Crystallographic data collection and structure determination

X-ray data was collected at beamline 23-ID-D of the Advanced Photon Source and processed with the program HKL2000(40). Phases from MR solutions were combined with those from SIRAS with the program Phenix(22). Cycles of rebuilding and refinement using the programs Coot(41, 42), Phenix(22), and Refmac(43) at *R*_work_ and *R*_free_ of 13.5% and 15.8%, respectively.

## Abbreviations

LRRK2: Leucine-Rich Repeat Kinase 2
ROC: Ras of complex proteins
COR: C-terminal of ROC

## ACKNOWLEDGMENTS

QQH acknowledges funding from the NIH (R01GM111639, R01GM115844) and the Michael J. Fox Foundation. YT acknowledges funding from the NIH (R01GM111695) and US National Science Foundation (MCB-1157688). This research was supported in part by the Intramural Research Program of the NIH, National Institute on Aging (MRC). SMJ acknowledges funding from the NIH (R01GM120350). NCH acknowledges funding from the Travel and Learn program of the Indiana Association of Chinese-Americans (IACA). GM/CA@APS has been funded in whole or in part with Federal funds from the National Cancer Institute (ACB-12002) and the National Institute of General Medical Sciences (AGM-12006). This research used resources of the Advanced Photon Source, a U.S. Department of Energy (DOE) Office of Science User Facility operated by Argonne National Laboratory under Contract No. DE-AC02-06CH11357.

## Author Contribution

C.W. and J.L. designed and performed experiments as well as prepared the manuscript; Y.P., V.A.E., L.W., and M.O. created DNA constructs and other research agents as well as assisted with manuscript preparation; N.C.H performed the molecular dynamics simulations; R.S. assisted with X-ray data collection and analysis; S.M.J. assisted experimental design and manuscript preparation; M.W. assisted data acquisition and analysis, as well as manuscript preparation; A.B., X.R., and M.R.C. provided reagents and assisted manuscript preparation; M.F., Y.T., A.B, and J.N. provided key research reagents; Q.Q.H. designed experiments, analyzed data, and prepared the manuscript.

## Structural Coordinates

Coordinates deposited to the Protein Data Bank (PDB ID: 6OJE and 6OJF).

