## Supplemental figures for "A revised 1.6 Å structure of the GTPase domain of the Parkinson’s disease-associated protein LRRK2 provides insights into mechanisms"

**Table 1 Data collection and refinement**

| | | | |
|---|---|---|---|
| **Data collection** | | | |
| Data set | $ROC_{KA}$ | $ROC_{WT}$ | $ROC_{WT}$-SeM |
| Space group | $P2_1$ | $P2_1$ | $P2_1$ |
| Unit cells | 44.63 101.88 44.61 | 44.65 103.69 44.59 | 44.58 102.95 44.61 |
| | 90.00 100.95 90.00 | 90.00 101.20 90.00 | 90.00 101.31 90.00 |
| Wavelength (Å) | 1.0331 | 1.0331 | 0.9794 |
| Resolution (Å)[1] | 43.82-1.59 | 43.8-1.88 | 51.47-3.00 |
| | (1.64-1.59) | (1.95-1.88) | (3.19-3.00) |
| Completeness (%) | 97.37 (78.93) | 97.41 (82.63) | 98.1 (97.4) |
| $R_{merge}$[2] | 0.084 (0.577) | 0.072 (0.457) | 0.064 (0.122) |
| $R_{meas}$[3] | 0.087 (0.623) | 0.078 (0.516) | 0.090 (0.172) |
| $R_{pim}$[4] | 0.023 (0.223) | 0.031 (0.233) | 0.064 (0.122) |
| $CC_{1/2}$ | (0.804) | (0.874) | (0.937) |
| I/σ(I) | 41.9 (1.7) | 24.0 (2.3) | 10.8 (6.2) |
| **Refinement** | | | |
| Unique reflection | 50680 (4069) | 31422 (2692) | |
| Protein atoms | 3029 | 2973 | |
| Solvent atoms | 235 | 237 | |
| Ligands | 60 | 60 | |
| R-factor ($R_{free}$) (%)[5] | 13.50 (15.76) | 18.75 (24.72) | |
| Average B-factor ($Å^2$) | 41.31 | 46.97 | |
| R.M.S. deviations | | | |
| Bonds (Å) | 0.023 | 0.019 | |
| Angles (°) | 2.62 | 2.26 | |
| **Ramachandran plot** | | | |
| most favored regions (%) | 89.19 | 95.07 | |
| Additionally allowed regions (%) | 7.3 | 3.84 | |
| Outlier regions (%) | 3.51 | 1.1 | |

[1]Values for the highest resolution shell are indicated in parentheses.

[2] $R_{merge} = \Sigma_h\Sigma_i |I_{hi}-\langle I_h\rangle|/\Sigma_h\Sigma_i I_h$, where $I_{hi}$ is the intensity of the $i^{th}$ observation of reflection h, and $\langle I_h\rangle$ is the average intensity of redundant measurements of the h reflections.

[3] $R_{meas} = \Sigma_h \sqrt{}(n/(n-1)\Sigma_i |I_{hi}-\langle I_h\rangle|/\Sigma_h\Sigma_i I_h$, where $I_{hi}$ is the intensity of the $i^{th}$ observation of reflection h, and $\langle I_h\rangle$ is the average intensity of redundant measurements of the h reflections.

[4] $R_{pim} = \Sigma_h \sqrt{}(1/(n-1)\Sigma_i |I_{hi}-\langle I_h\rangle|/\Sigma_h\Sigma_i I_h$, where $I_{hi}$ is the intensity of the $i^{th}$ observation of reflection h, and $\langle I_h\rangle$ is the average intensity of redundant measurements of the h reflections.

[5]R-factor = $\Sigma||F_o|-|F_c||/\Sigma|F_o|$, where $F_o$ and $F_c$ are the observed and calculated structure-factor amplitudes. $R_{free}$ is monitored with 5% of reflections excluded from refinement.

**Figure S1. Structure and activity of WT vs. KA surface engineered ROC$_{ext}$. a**) Superposition of WT (gold) and a surface-engineered (K1460A, K1463A) construct of ROCext (green), showing that the two structures are practically the same. **b**) GTPase activity of the surfaced engineered mutant (KA, red), WT (blue), and a PD-associated mutant R1441H (green); showing that the surface engineering did not significantly affect its GTPase activity compared to the WT.

a

b

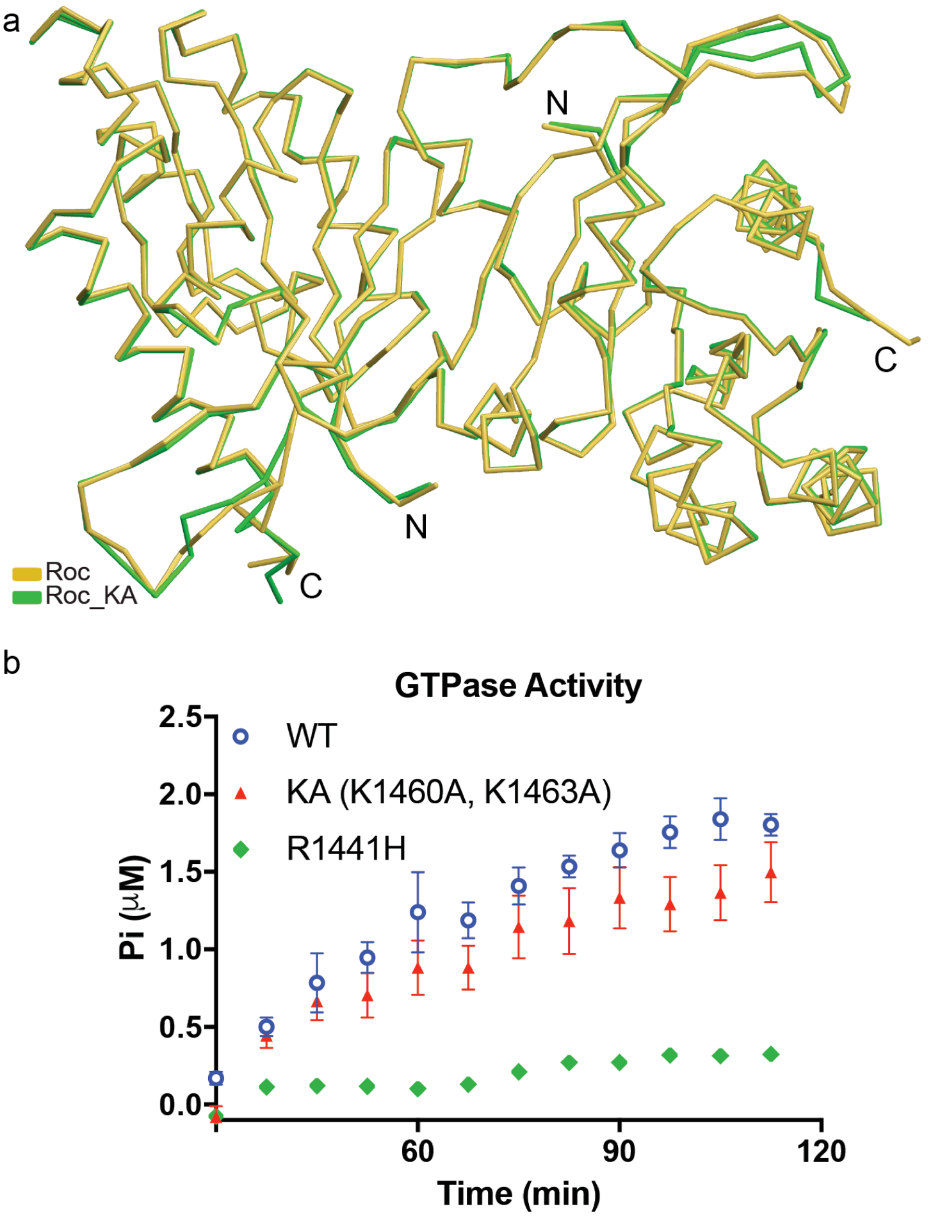

**Figure S2. Backbone tracing of ROC$_{ext}$ and 2ZEJ**. Backbone tracing and electron-density (2FoFc map contoured at 1σ) of the 2ZEJ structure (left panel) and ROC$_{ext}$ (right panel) showing a lack of density after residue K1356 in the map of 2ZEJ (blue mesh), but are clearly defined in the map of ROC$_{ext}$ (magenta mesh).

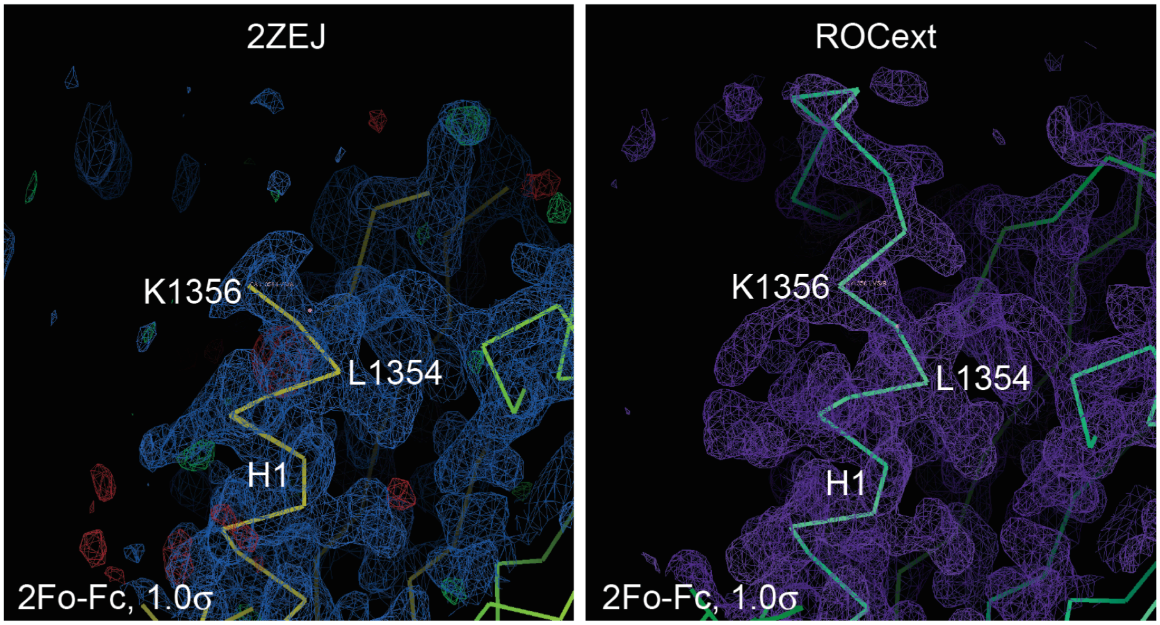


**Figure S3. Backbone tracing of ROC$_{ext}$ and 2ZEJ**. Backbone tracing and electron-density (2FoFc at 1σ) of the 2ZEJ structure (left panel) and ROC$_{ext}$ (right panel) showing the 2ZEJ tracing is discontinuous at T1368, but the same region is clearly defined in the map of ROC$_{ext}$. Note that the 2ZEJ map shows positive difference (Fo-Fc) density (green) that tracks with the tracing of ROC$_{ext}$ (SW1), which is evident that the unbuilt SW1 of 2ZEJ is located at the same location as that of ROC$_{ext}$.

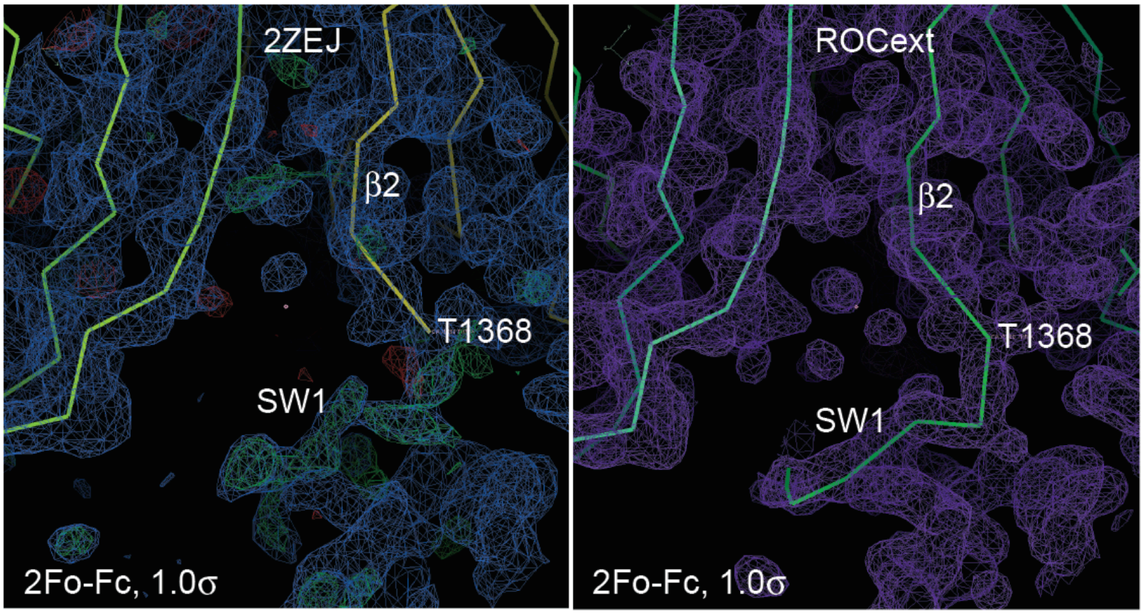

**Figure S4. 2ZEJ structure with the chain assignment of H1 and β1 swapped**. **a**) and **b**) superposition of $ROC_{ext}$ (gold) with 2ZEJ (blue) after swapping the chain assignment of H1 and β1 from chain A to chain B and vice versa, showing that the two structures are practically the same after the chain-swapping.

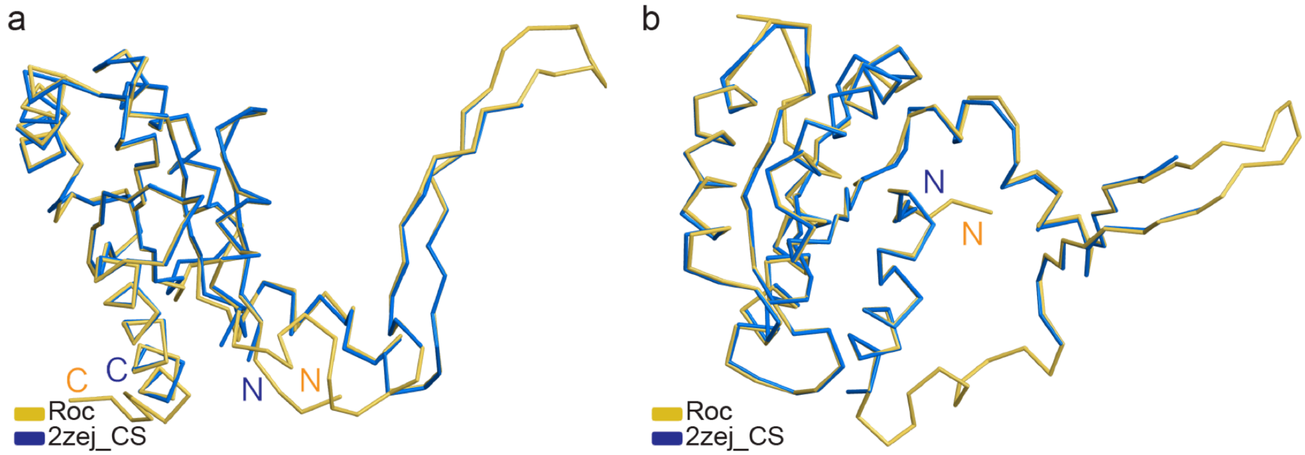


**Figure S5. Molecular simulation of disulfide-stabilized $ROC_{ext}$. a**) Ribbon presentation of a molecular dynamic simulation model of ROC dimer consisting of an engineered disulfide bond between residues 1398 and 1431 (S-S). **b**) Crystal structure of $ROC_{ext}$ (light grey) superimposed with the calculated S-S dimer (gold and teal) (RMSD 0.93 $A^2$), showing no significant structural changes upon disulfide formation. The engineered disulfide bonds are highlighted with dotted lines.

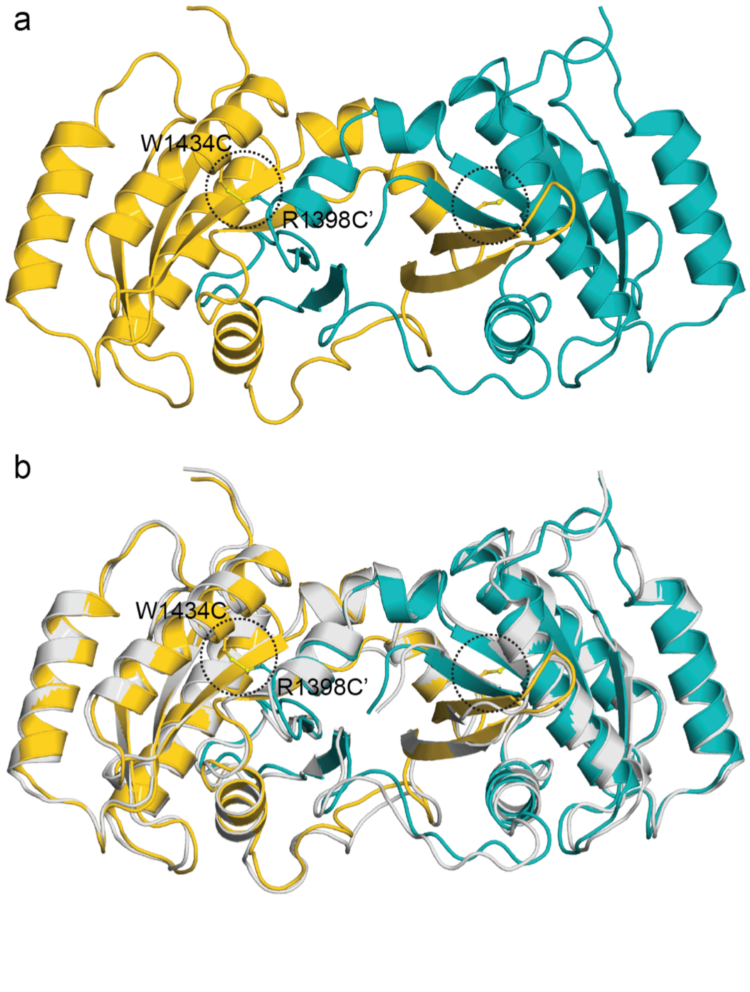

**Figure S6. Disulfide-stabilized dimer**. Size-exclusion chromatography coupled with multi-angle light scattering, showing that the elution profile and calculated mass of the disulfide-stabilized dimer (dashed line) was comparable to wild-type dimer (solid black line).

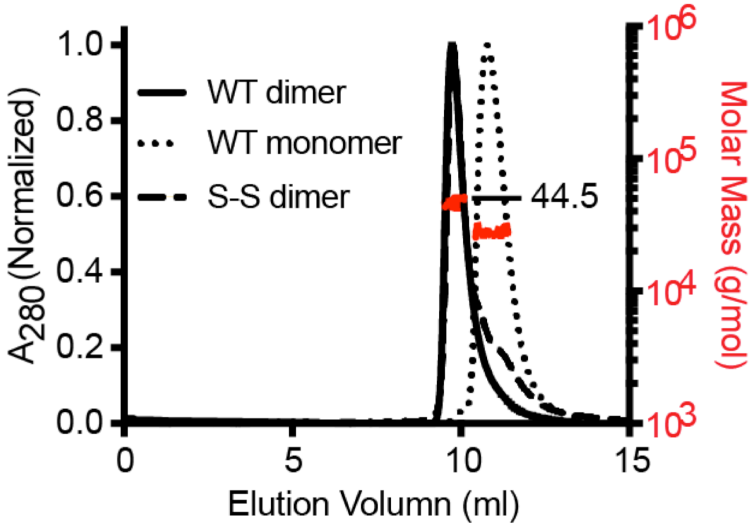


**Figure S7. SDS-PAGE of disulfide-stabilized dimers and monomers**. **a**) Non-reducing SDS-PAGE showing the S-S monomer (S1) migrating at about 22 kDa and the S-S dimer (S2) at about 44 kDa. **b**) Reducing SDS-PAGE showing that both monomers (S1) and dimers (S2) migrating at about 22 kDa.

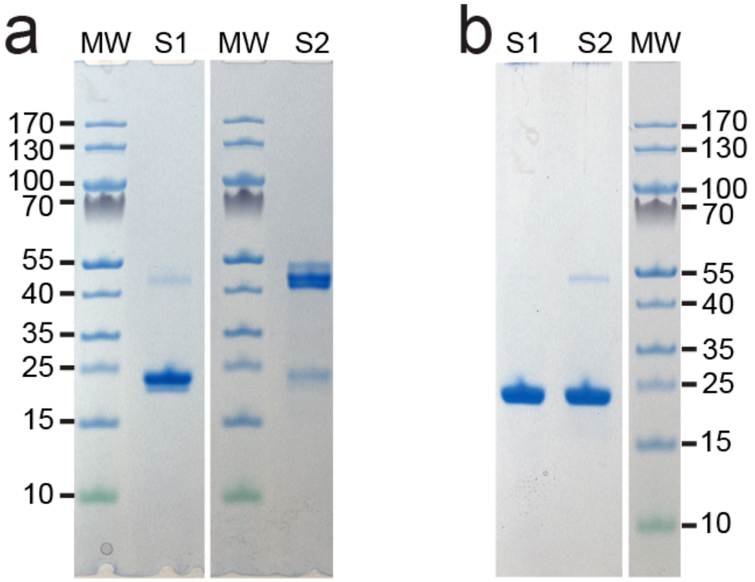

**Figure S8. γ-Phosphate interacting residues**. Superposition of GTP-bound conformation of Ras with a monomeric model of $ROC_{ext}$ showing residues Thr1368 and Gly1397 are in the same locations as Thr35 and Glyl60 of Ras.

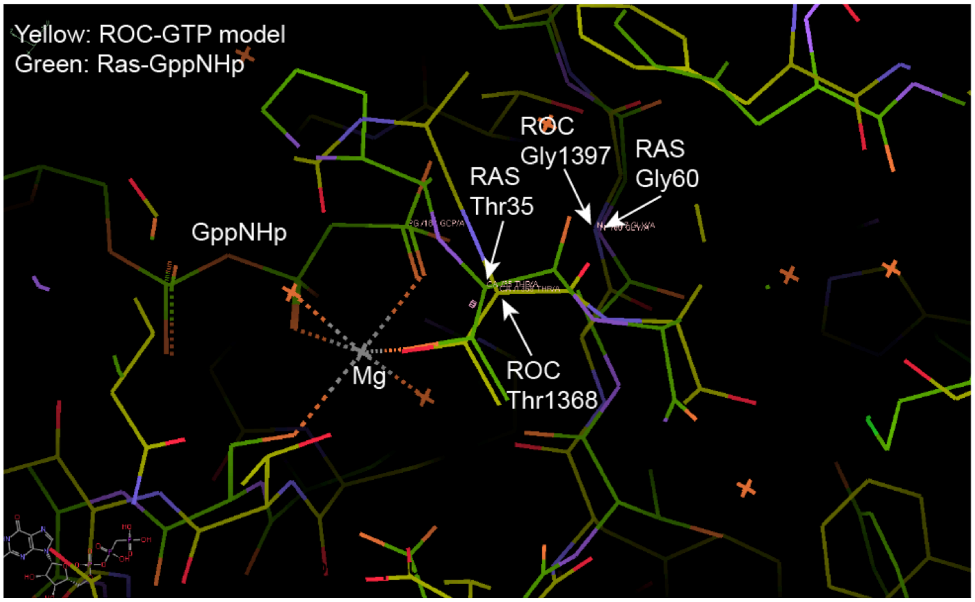


**Figure S9. Interactions at residue R1441**. Enlarged view of the $ROC_{ext}$ dimer interface around residue R1441, showing the surface complementarity between the sidechain of R1441 with the concave surface around residue F1401. Chain A is shown as yellow ribbons and chain B is shown as red surface rendering. Residue R1441 is shown as rod-and-ball model.

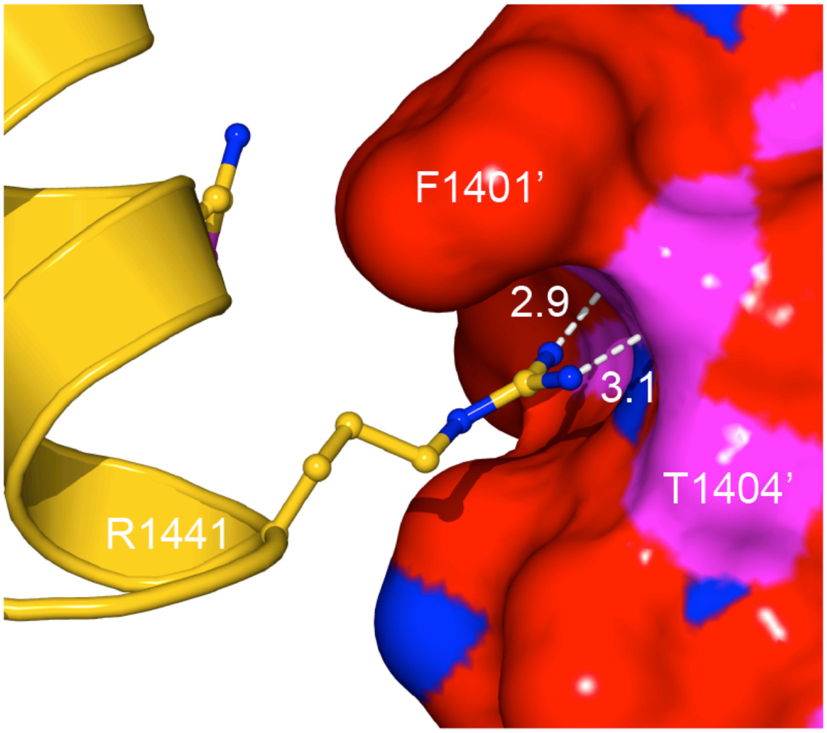

**Figure S10. Crystals of WT and the surface engineered KA construct**. Optimized crystals of WT $ROC_{ext}$ (left panel) and the surface engineered K1460A-K1463A (right panel), showing improvement of crystals from clusters of thin plates (left panel) to a single crystal with increased thickness along the b-axis (right panel).

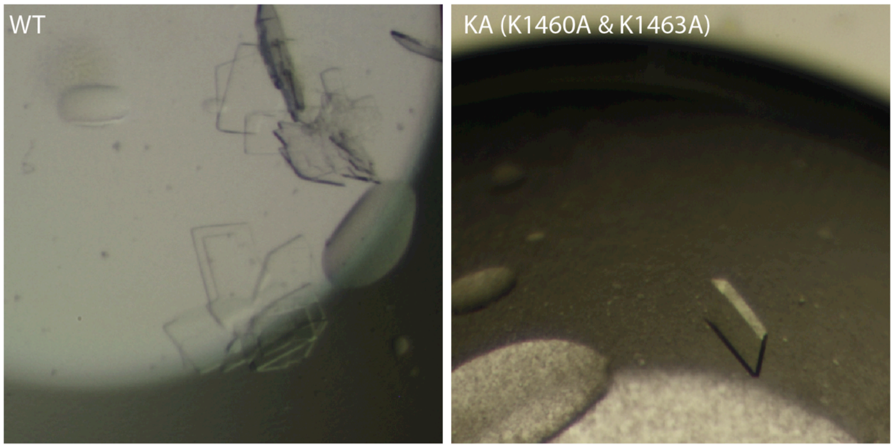

### Methods

#### Protein Expression and Purification

An extended GTPase domain of LRRK2 consisting of residues 1329-1520 ($ROC_{ext}$) was subcloned into a pETDuet-1 vector (Novagen, Merch KgaA, Darmstadt, Germany) using PCR cloning techniques. The resulting protein consisting of an N-terminal hexahistidine tag was expressed from Rosetta2 (DE3) *E. coli* (Novagen) by inducing with 0.5 mM isopropyl-β-D-thiogalactopyranoside (IPTG) for 16 hrs at 20 °C. Cells were harvested by centrifugation and lysed by sonication in a buffer containing 30 mM HEPES (pH 7.4), 250 mM NaCl, 10 mM $MgCl_2$, 10 mM Glycine, 20 mM imidazole, 10 $\mu$M GDP, and 10% (v/v) Glycerol. Cell debris was cleared by ultra-centrifugation at 14,000 g (35,000 rpm, Beckman 45 Ti rotor). The supernatant was incubated with Ni-NTA Agarose (Invitrogen) for 2 hrs at 4 °C, then washed with lysis buffer (detailed above) and eluted with buffer containing 30 mM HEPES (pH 7.4), 250 mM NaCl, 10 mM $MgCl_2$, 10 mM Glycine, 300 mM imidazole, 1 mM DTT, 10 $\mu$M GDP, and 10% Glycerol. The purified protein was then 'polished' by passing through a size-exclusion column (Superdex 200, GE Healthcare) in buffer containing 30 mM HEPES pH 7.4, 150 mM NaCl, 10 mM $MgCl_2$, 10 mM Glycine, 1 mM DTT, and 10% Glycerol. The purified protein was then concentrated to ~15 mg/mL, flash frozen in liquid nitrogen and stored at -80 °C. The disulfide bond-stabilized ROC dimer (S-S), consisting of R1398C and W1434C double mutation, was subcloned in the pETDuet-1 vector and expressed from SHuffle T7 Express *lysY E. coli* (New England Biolabs Inc. MA, USA) by inducing with 0.5mM IPTG for 16 hrs at 20°C. The purification of the ROC S-S was performed as described for the wild-type.

#### Expression and Purification of Selenomethionine Substituted Protein

SeMet $ROC_{ext}$ was expressed from Rosetta2 (DE3) *E. coli* (Novagen). Rosetta2 (DE3) cells were grown in the M9 minimal medium. Methionine synthesis was inhibited by adding 100 mg of Lys, Phe, Thr (Sigma Aldrich) and 50 mg of Ile, Leu, Val (Sigma Aldrich) per liter of M9 minimal medium. At the OD of 0.6 – 0.8, 60 mg of L-Selenomethionine (Sigma Aldrich) was supplied to the medium while cells were induced by addition of 0.5 mM IPTG at 20°C for overnight. The SeMet $ROC_{ext}$ purification was the same as wild-type $ROC_{ext}$.

#### Size-Exclusion Chromatography Coupled with Multi-angle Light Scattering (SEC-MALS)

To determine the absolute molecular weight of $Roc_{ext}$ in solution, we used multiple angle light scattering. Our experimental setup includes an AKTA FPLC (GE Healthcare Biosciences, Piscataway, New Jersey) with a silica-based size-exclusion chromatography column (WTC-030S5, Wyatt Technology Corporation, Santa Barbara, California) as a liquid chromatography (LC) unit. Down from the LC is a refractive index detector (Optilab T-rEX, Wyatt Tech.) followed by a multiple light scattering detector (Dawn HeleosII, Wyatt Tech.) for determining protein concentration and particle size, respectively. Each sample injection consisted of ~1 mg of purified $ROC_{ext}$ in buffer containing 30 mM HEPES (pH 7.4), 0.15 M NaCl, 10 mM $MgCl_2$, 10 mM Glycine, 1 mM DTT, and 10% Glycerol. The flow rate was set at 0.4 mL/min and data were collected in a 1-second interval. Data processing and analysis were performed using the ASTRA software (Wyatt Tech.)

#### Circular Dichroism Spectroscopy

CD spectra were collected on a Biologic Science Instruments MOS450 AF/CD spectrometer with a slit width of 1.0 mm and data acquisition of 1.0 s. The protein samples with concentrations ranging 0.46 - 0.86 mg/mL (based on absorbance at 280 nm) were dissolved in the buffer containing 10 mM Tris-HCl (pH 7.4), 150 mM NaCl, 5 mM $MgCl_2$, 1 mM DTT, and 5% Glycerol.

#### Fluorescence Polarization Nucleotide-binding Assay

To estimate the binding affinity of guanine nucleotides BODIPY-FL-GTPgammaS (100 nM) or BODIPY-FL-GDP (150 nM) (Molecular Probes) were titrated with $ROC_{ext}$ (starting at 0.1 $\mu$M) until saturation was reached (15 $\mu$M and 10 $\mu$M, respectively). Fluorescence polarization signals were read using an EnVison 2102 Multilabel Plate Reader (Perkin Elmer, Massachusetts) with excitation at 485 nm and emission at 535 nm. Experiments were performed at 25 °C in buffer containing 30 mM HEPES (pH 7.4), 150 mM NaCl, 10 mM $MgCl_2$, 10 mM Glycine, 4 mM EDTA, 1 mM DTT, and 10% Glycerol. Data were analyzed using Prism 6 (GraphPad Software, CA, USA).

#### GTPase Activity Assay

GTPase activity of $ROC_{ext}$ was assessed by using the Enzcheck assay kit (Invitrogen) according to the manufacturer's instructions. Briefly, $Roc_{ext}$ (30 μM) was incubated with 2 mM GTP in buffer containing 30 mM HEPES (pH 7.4), 150 mM NaCl, 10 mM $MgCl_2$, 10 mM Glycine, 1 mM DTT, and 10% Glycerol at 25 °C. Absorbance at 360 nm was recorded in every 3 minutes for 3 hours using a microplate reader. The amount of inorganic phosphate released from GTP hydrolysis at each time points was determined by extrapolation using a phosphate standard curve. In experiments comparing the activity of the S-S mutant with WT, DTT was omitted from the reaction for both the S-S mutant and WT. Data analysis and curve fitting were done with GraphPad Prism 7.

#### Thermofluor Assay

Solutions of 12.5 μL of 10x Sypro Orange (prepared from 5,000x stock concentrate, Molecular Probes) in buffer containing 30 mM HEPES pH 7.4, 0.15 M NaCl, 10 mM $MgCl_2$, 10 mM Glycine, 1 mM DTT, 4 mM GDP or Gpp(NH)p, 2 μM LMNG, and 10% Glycerol, and 12.5 μL of 25 μM $ROC_{ext}$ or mutants were added to a 96-well thin-wall PCR plate. The plate was heated in the Real-Time PCR Detection System (Mastercycler realplex, Eppendorf) from 20 to 85 °C and fluorescence recorded in increments of 0.4 °C. The emission wavelength was set at 550 nm.

#### Homology Modeling and Molecular Simulation

Homology models of the monomeric ROC domain in GDP-/GTP-bound and apo conformations were built based on the structures of Ras and *C. tepidium* Roco (PDB ID: 4Q21, 6Q21, 3DPU, and 6HLU, respectively) by using the program Modeller 9.19 (Andrej Sali, UCSF, San

Francisco, CA). Molecular simulation of the S-S mutant was performed using CHARMM[38] and NAMD[39], and Molecular graphics display and presentation were made using PyMol (www.pymol.org).

#### Crystallographic data collection and structure determination

A 1.6 Å data set from crystals of $ROC_{ext}$ was collected at beamline 23-ID-D of the Advanced Photon Source at Argonne National Laboratory. The X-ray wavelength was set at 1.03 Å. We cryo-cooled $ROC_{ext}$ crystals in liquid nitrogen after transferring them to a cryo-protectant containing 10% glycerol, 100 mM KSCN, 25% PEG MME, and 0.1M BisTris, pH 6.5. Diffraction data were collected at 100 K on a Pilatus3 6M detector (DECTRIS, Switzerland) and processed with the program HKL2000[40]. The crystals belonged to the space group $P2_1$ with two molecules per asymmetric unit. The homology model was used as a molecular-replacement search model in combination with single anomalous dispersion (SAD) and yielded a solution with the program Phenix[22]. A 2fo-2fc map following rigid body refinement showed reasonable density for ~80% of the model. Data of Se-Met $ROC_{ext}$ crystals to 3.0 Å resolution were collected on the same beamline. The X-ray wavelength was set at 0.98 Å. Cycles of rebuilding and refinement were carried out using the programs Coot[41,42], Phenix[22], and Refmac[43]. The current $R_{cryst}$ and $R_{free}$ are 13.5% and 15.8%, respectively (r.m.s. deviations from ideal bond lengths and angles are 0.023 Å and 2.6°, respectively).
